# First Successful Targeted Mutagenesis Using CRISPR/Cas9 in Stably Transformed Grain Amaranth Tissue

**DOI:** 10.1101/2025.02.22.639339

**Authors:** Susanne K Vollmer, Markus G Stetter, Götz Hensel

**Author notes:** **Email addresses:**.

## Abstract

Genome editing using CRISPR/Cas is a key technology for speeding up breeding for climate-resilient, high-yielding crops (Scheben *et al*., 2017). However, efficient targeted mutagenesis requires implementing stable transformation methods and establishing a CRISPR/Cas setup suitable for the species of interest (Shan *et al*., 2020). The availability of such methods is a significant bottleneck to advancing many promising, albeit under-researched, crops. Testing and establishing vectors for efficient application of CRISPR/Cas in non-model crops could boost research and breeding of new valuable crops (Ye and Fan, 2021).

We edited key pathway genes in the betalain biosynthesis pathway of grain amaranth, i.e., *A. hypochondriacus L*., to prove how targeted mutagenesis can be implemented in an orphan crop using the CasCADE modular cloning system (Hoffie, 2022). Grain amaranth is a resilient C_4_ dicot orphan crop with excellent nutritional composition. These properties make amaranth a well-suited candidate to be bred as a climate-resilient crop (Joshi *et al*., 2018). However, no efficient and reproducible protocol for successful application of CRISPR/Cas9 or stable transformation and regeneration, has been demonstrated in *A. hypochondriacus* (Castellanos-Arévalo *et al*., 2020).

## Main part

Amaranth produces red and yellow betalains, specialized metabolites in *Caryophyllales* species (Timoneda *et al*., 2019). Betalains have been employed as reporters in molecular biology using the RUBY cassette, which consists of the three enzymes required to produce red betalains from tyrosine (He *et al*., 2020). Conversely, the pathway is well suited to evaluate the knock-out efficiency through targeted mutagenesis in *A. hypochondriacus*, where the pigments naturally occur. Key enzyme genes in the betalain pathway are *AhCYP76AD2* and *AhCYP76AD5*, which catalyze the initial steps of betalain production. Natural knock-outs of *AhCYP76AD2* have been shown to lack red color (Winkler *et al*., 2024), suggesting its suitability as a visual reporter.

The foundations of successful CRISPR/Cas-mediated targeted mutagenesis are efficient guide RNAs and suitable components to express the Cas9 enzyme and the gRNAs (Shan *et al*., 2020). The *Cas9* sequence should be adapted to the codon usage of the target species. The promoter to drive the expression of *Cas9* and the guides need to be vigorously active in the target species. Finally, the selection cassette has to function efficiently in the target species. To address these requirements in amaranth, we constructed a binary vector containing Cas9 and a four-guide cassette, using the CasCADE modular cloning system (**Figure 1a, Supplemental Figure 1, Supplemental Table 1, Appendix 1;** Hoffie, 2022). Two guides were designed to target the first exon of *AhCYP76AD2* (gRNA 1 and gRNA 2) and one to target the first exon of *AhCYP76AD5* (gRNA 4, **Figure 1a**). Amaranth plants successfully mutated in *AhCYP76AD2* should be deficient in betalains, facilitating a later detection of successfully edited regenerates. An additional gRNA not targeting the amaranth genome, but the *CYP76AD2* gene in the RUBY reporter cassette (He *et al*., 2020; gRNA 3) was included as control.

**Figure 1:**
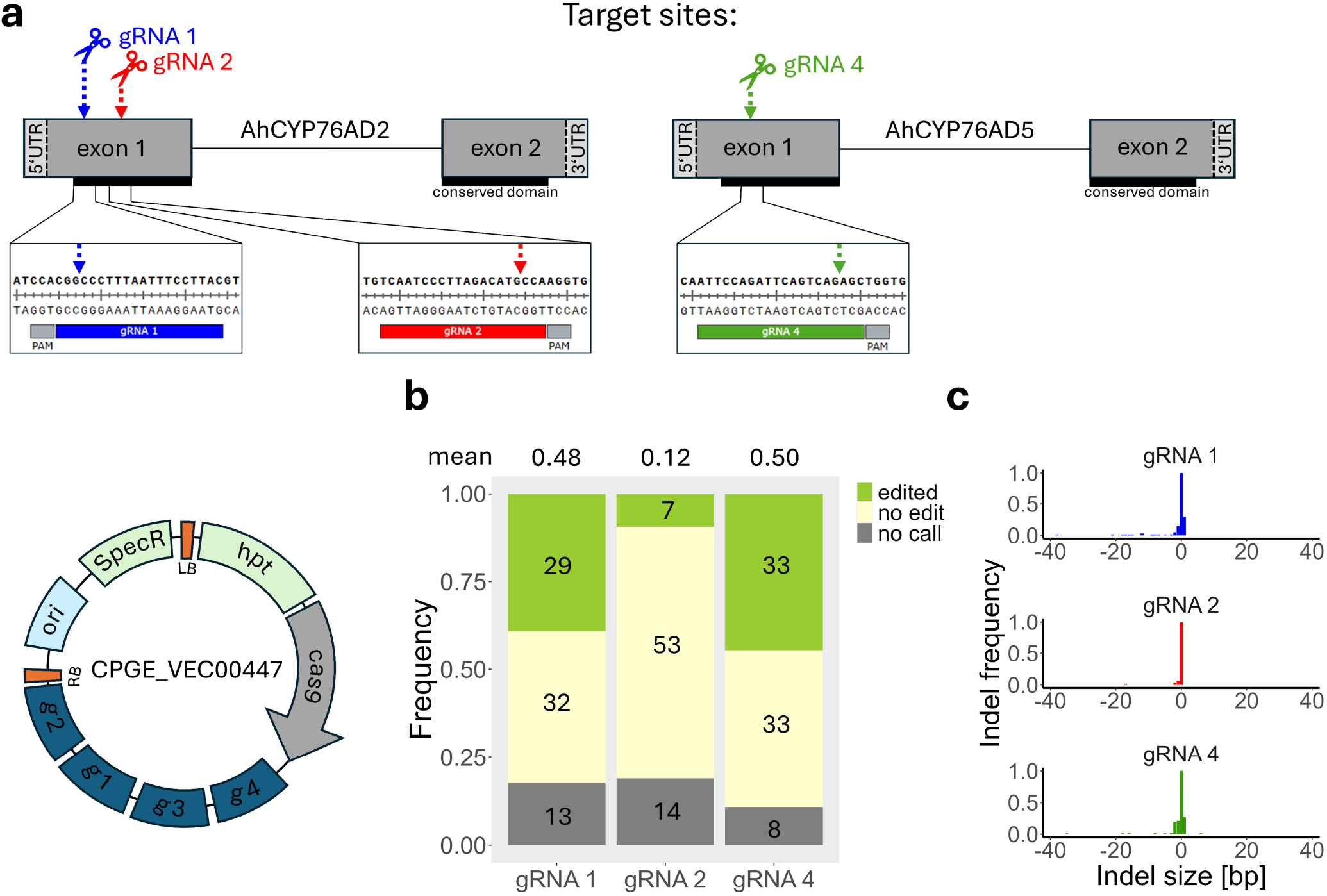
Overview of CRISPR/Cas9-mediated targeted mutagenesis in grain amaranth. a) Vector with four gRNA arrays, two targeting the first exon of *AhCYP76AD2* (gRNA 1 and gRNA 2), one the first exon of *AhCYP76AD5* (gRNA 4) and one the *RUBY* reporter (gRNA 3). Schematic structure of the target genes with the respective target sites of the guides in amaranth genes. b) Total number of mutations for all three guides. c) Length and frequency of the observed indels found in 74 calli.

The *Cas9* gene was codon-optimized for the dicot *Arabidopsis thaliana* and driven by the parsley *Ubi4-2* promoter (Fauser *et al*., 2014; Hoffie, 2022). Each gRNA was expressed separately by the *AtU6-26* promoter (Hoffie, 2022). For the transgenic tissue selection, the T-DNA contains an intronized *hpt* gene driven by a *CaMV* doubled-enhanced *35S* promoter to confer hygromycin resistance (Hoffie, 2022). We cloned the CRISPR/Cas9 cassette into a binary vector via SfiI to enable *Rhizobium*-mediated transformation (Hoffie, 2022).

Next, we established callus transformation, which enables the highly efficient production of stable transgenic tissue in multiple-grain amaranth species. The results were obtained from two independent transformation replicates, batch 1 and 2. A random subset of 10 calli from batch 1 and all from batch 2 treated with *R. radiobacter* carrying the CRISPR/Cas9 vector were analyzed for edits. Briefly, 5-week-old calli of *A. hypochondriacus* were transformed with the CRISPR/Cas9 vector using *R. radiobacter* strain GV3101. For co-culture, calli were incubated on filter paper for three days before being transferred to a callus induction medium with selection. All calli treated with *Rhizobium* carrying the CRISPR/Cas9 vector (42/42 for batch 1 and 65/65 for batch 2) grew new resistant calli on the selection medium.

In contrast, only two (1/21 for batch 1 and 1/32 for batch 2) of the non-treated calli survived the selection and propagated new callus, indicating the high success of transformation. After multiple rounds of selection, callus material was sampled for genotyping and analysis (**Supplemental Figure 2**). Genomic DNA (gDNA) was extracted from 74 calli of the two independent batches.

We sampled individual calli; however, callus tissue can contain cells from independent transformation events with different edits. Therefore, each gDNA included a pool of wild-type and edited alleles. To increase the sensitivity for detecting edited alleles using Sanger sequencing, we employed restriction enzyme digestion-suppressed PCR (RE-PCR) for two guides (gRNA 1 and gRNA 4) in the amaranth genome. Through the deconvolution of the obtained chromatograms, we found edits in 50% (gRNA 4) and 48% (gRNA 1) of the analyzed calli (**Figure 1b**). In contrast, for gRNA 2, a lack of a restriction enzyme recognition site at the Cas9-mediated cut site precluded RE-PCR, edits could only be detected in 12% of the samples. Among all samples with data available for the three sites, 26.8% were edited for both target sites, and 7.1% for all three guide positions.

To ensure complete knockouts for downstream analysis of the target gene, targeted mutagenesis should achieve a diverse set of mutations including larger indels. According to the deconvolution data, most edits resulted in a one-base pair deletion or insertion (**Figure 1c**). However, for each guide, edits with larger deletions were also found (max 35 bp for gRNA 4, 17 bp for gRNA 2, 38 bp for gRNA 1; **Figure S3**). To confirm the results from the deconvolution, we cloned the target region of gRNA 4 and gRNAs 1 and 2 from a subset of samples into pGEM-Teasy (Promega) and sequenced multiple colonies. While the frequency of mutations differed, the types of mutations mostly agreed across both methods (despite the restricted sample size of eight sequenced clones from the subcloning) among samples (**Supplemental Table 2**).

Our results show the first successful targeted mutagenesis in grain amaranth using CRISPR/Cas9. Using the CasCADE modular cloning system and our established callus transformation, we edited approximately 50% of all analyzed calli for both target genes. This paves the way for genome editing in grain amaranth for research and breeding of this orphan crop. Moreover, it may also guide the design of CRISPR systems for other species of the *Caryophyllales*, where reports of successful edits are still scarce.

## Acknowledgements

The Deutsche Forschungsgemeinschaft supported this work under Germany’s Excellence Strategy – EXC-2048/1 (project ID 390686111), and STE 2654/4.

## Author Contributions

M.G.S. and G.H. supervised the project. S.K.V. performed the experiments and analyzed the data. S.K.V., M.G.S. and G.H. discussed and interpreted the data. S.K.V. wrote the manuscript and prepared the figures. All authors edited and approved the manuscript.

## Supporting Information

**Supplemental Figure 1**– Plasmid map of vector used for the targeted mutagenesis (CPGE_VEC00447)

**Supplemental Figure 2**– Genotyping of calli for the presence of the *Cas9* gene **Supplemental Figure 3**-Gel electrophoresis of 35 bp deletion at target site of gRNA 4 in one sample

**Supplemental Table 1**– Primers used in this study

**Supplemental Table 2**– Comparison of editing frequency from subcloning and deconvolution

**Appendix S1** – DNA sequence of the vector (CPGE_VEC00447)

