## Supplementary material for "First Successful Targeted Mutagenesis Using CRISPR/Cas9 in Stably Transformed Grain Amaranth Tissue": Suppl methods and figures

Susanne K Vollmer^1,2,3^, Markus G Stetter^2,3^ and Götz Hensel^1,3*
*^ corresponding author

**Supplemental methods**

**S1 Cloning of binary plasmid CPGE_VEC00447 for targeted mutagenesis**

Cloning was conducted using the CasCADE modular cloning set (Hoffie, 2022). Target sequences (Supplemental Table 2) were designed with the CRISPOR (<https://crispor.gi.ucsc.edu/>), CRISPRscan (<https://www.crisprscan.org/>), and RNAfold (<http://rna.tbi.univie.ac.at/cgi-bin/RNAWebSuite/RNAfold.cgi>) webtools. The chosen target sequences were ordered with the necessary overhangs for Golden Gate cloning using the BbsI Type IIS restriction enzyme as oligos (Biolegio B.V., Nijmegen, The Netherlands). A 5 µl mixture of each respective forward and reverse oligo (10 µM) was prepared and incubated for 5 minutes at 95°C. For cloning into Level 0 vectors (IK75-IK78) harbouring the Arabidopsis U6-26 promoter and the Cas9-specific scaffold, 5 µl of the respective duplex and 1 µl of 30 ng/µl Level 0 plasmid were combined with 1 µl BbsI, 1 µl T4 ligase, 2 µl of 10xFD buffer, 1 µl ATP (10 nM), and 9 µl of milliQ water. Golden Gate cloning was executed in 10 cycles of 5 minutes at 37°C and 10 minutes at 22°C, followed by 30 minutes at 37°C and 15 minutes at 75°C. Five µl of the Golden Gate reactions were used to transform *E. coli* Mach1 or XL1 cells, generating plasmids CPGE_VEC00441-CPGE_VEC00444. For Level 1 cloning, 150 ng of each Level 0 vector and 30 ng of the Level 1 backbone (IK19) were utilised for Golden Gate cloning with Esp3I, generating vector CPGE_VEC00445. For Level 2 cloning, each 150 ng of the gRNA assembly (CPGE_VEC00445), the Arabidopsis codon-optimised Cas9 expression cassette (RB20), the auxiliary unit (IK155), and the Level 2 backbone (IK48) were combined for Golden Gate cloning using BsaI generating CPGE_VEC00446. For cloning of the final binary vector, 20 ng of gel-purified SfiI-digested backbone (CPGE_VEC00113) and 60 ng of SfiI-digested Level 2 vector (CPGE_VEC00446) were mixed with 2 µl of T4 ligase buffer, 1 µl of T4 ligase, and 13 µl of milliQ water leading to CPGE_VEC00447. Successful cloning was verified via colony PCR, test digest (excluding Level 0 vectors), and Sanger sequencing (Microsynth, Balgach, Switzerland). The final CPGE_VEC00447 was further examined through whole plasmid sequencing (Microsynth, Balgach, Switzerland).

**S2 Callus transformation**

Seeds of *A. hypochondriacus* (PI 558499) were sterilized (2 min in 70% ethanol, 20 min in 2.4% sodium hypochlorite with one drop of Tween-20, five washes with sterile water) and germinated on germination medium (1/2 MS + 20 g/l sucrose, pH5.8, 8 g/l agar) at 24 °C dark to cut hypocotyl explants after 3 days. Explants were cultivated on callus induction medium (full MS (with vitamins) + 2 mg/L 2.4D + 0.5 mg/l BAP + 20 g Sucrose, pH 5.8, 8 g/L agar) (Castellanos-Arévalo et al., 2020; Wang et al., 2001) for approximately 34-36 days at 24°C in the dark, with subculturing every 2-3 weeks, until a callus of at least 2 mm had formed. For transformation, *A. tumefaciens* GV3101 was grown in CPY medium without antibiotics and with 200 µm acetosyringone and OD_600_ adjusted to 0.7. Callus were incubated in the bacterial solution for 10 minutes at 50 rpm, then dried on sterile filter paper for 15 minutes. Thereafter, calli were transferred to fresh sterile filter paper on transformation plates (callus induction medium + 100 µM acetosyringone) and co-cultured for 3 days at 21°C in the dark. Without washing, explants were transferred to callus induction medium with 150 mg/l timentin and 20 mg/l hygromycin for selection and cultivated at 24°C in the dark. On average, explants were transferred to fresh plates every two weeks.

**S3 Genotyping**

For analysis, callus samples were flash frozen in liquid N_2_, ground with a pestle and mortar, and gDNA was extracted using the NucleoSpin Plant II kit (Macherey-Nagel, Düren, Germany). For RE-PCR, 12 µl of the extracted gDNA was incubated with 0.4 µl of enzyme (Hpy188I for g4 and HaeIII for g1), 2 µl of 10x cutsmart, and 5.6 µl of milliQ water. Subsequently, 0.75 µl of the digested gDNA was used in a 30 µl PCR reaction with 15 µl of 2x GoTaq MasterMix (Promega, Madison, Wisconsin, United States of America) and 1.5 µl of respective forward and reverse primer (10 µM). For amplification, the reaction was incubated for 5 minutes at 95°C, followed by 35 cycles of 95°C for 20 seconds, 58°C for 15 seconds, and 72°C for 1 minute, with a final extension of 7 minutes at 72°C. The purified PCR product was then sent for Sanger sequencing (Microsynth, Balgach, Switzerland) or used for subcloning in pGEM-Teasy (Promega, Madison, Wisconsin, United States of America).

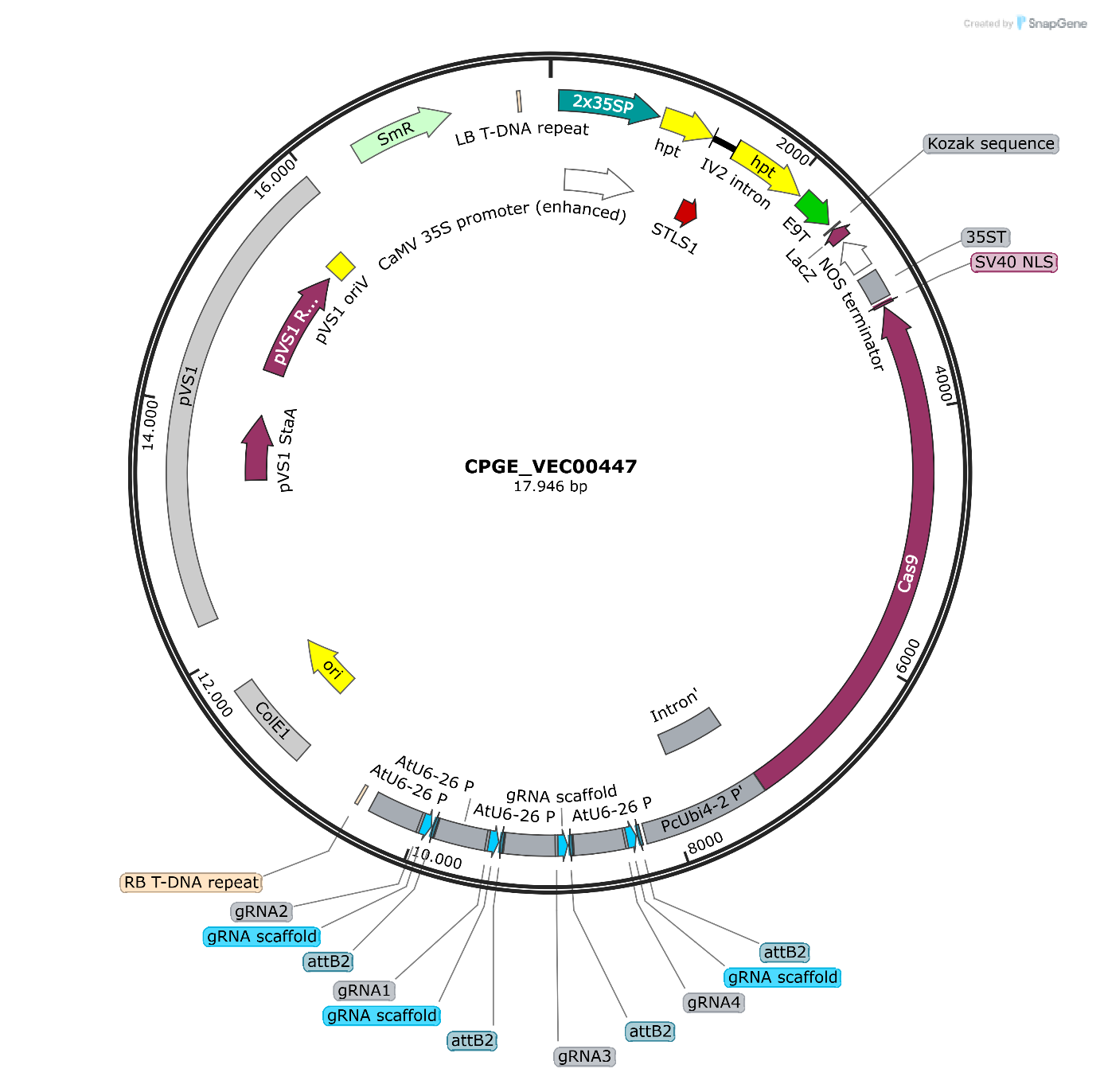
 **Supplemental Figure 1:** Plasmid map of the vector used for the targeted mutagenesis (CPGE_VEC00447).

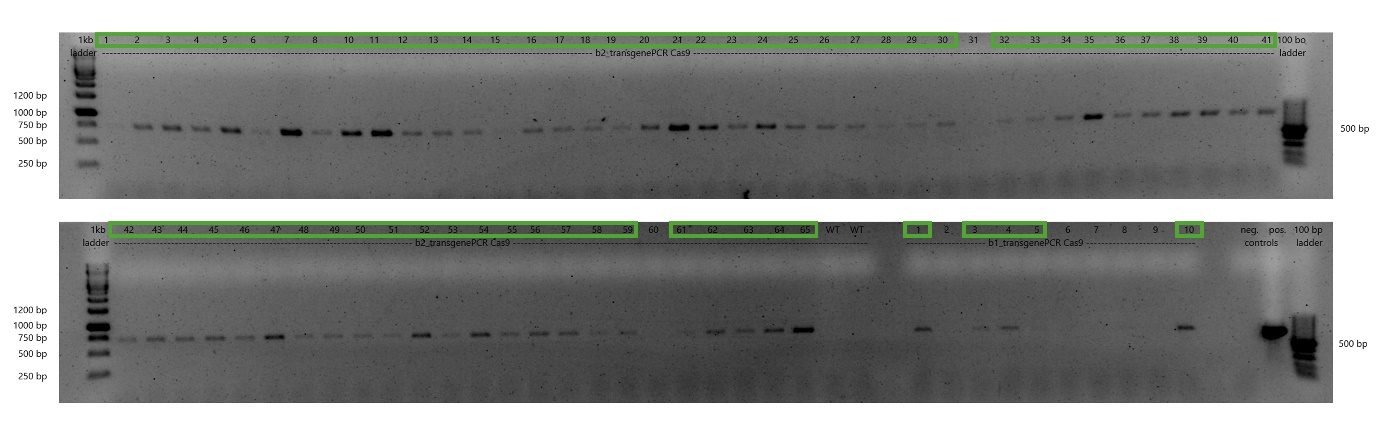
 **Supplemental Figure 2:** Genotyping of all 74 calli for the presence of the *Cas9* gene using primer pairs mentioned in Suppl Table 1. Samples with a visible amplification are marked with green (67/74).

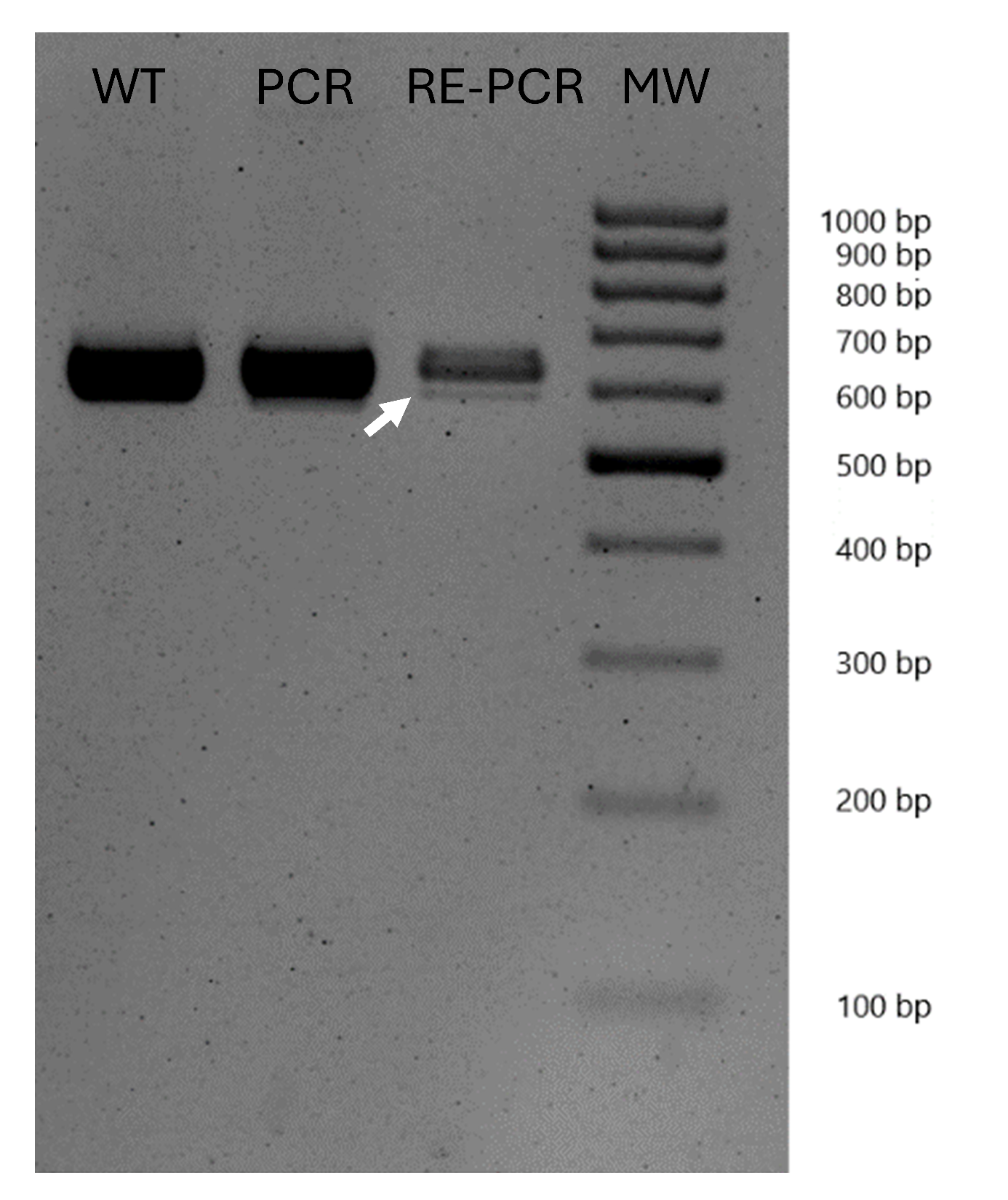
 **Supplemental Figure 3:** Gel electrophoresis of RE-PCR assay of sample with a 35 bp deletion at target site of gRNA 4.

**Supplemental Table 1:** Primers used for cloning and genotyping.

| Primer | Sequence (5‘- 3‘) | Purpose |
| --- | --- | --- |
| CPGE_PRM00523 | ATTGgcaatcccttagacatgcca | Forward and reverse primer for gRNA 2 sequence with overhangs |
| CPGE_PRM00524 | AAACtggcatgtctaagggattgc |  |
| CPGE_PRM00529 | AAACcggccctttaatttccttat | Forward and reverse primer for gRNA 1 sequence with overhangs |
| CPGE_PRM00530 | ATTGataaggaaattaaagggccg |  |
| CPGE_PRM00535 | AAACccgaccgtgttcttgctccc | Forward and reverse primer for gRNA 3 sequence with overhangs |
| CPGE_PRM00536 | ATTGgggagcaagaacacggtcgg |  |
| CPGE_PRM00541 | ATTGattccagattcagtcagagc | Forward and reverse primer for gRNA 4 sequence with overhangs |
| CPGE_PRM00542 | AAACgctctgactgaatctggaat |  |
| CPGE_PRM00531 | tcatggataatgcaaccctagc | Amplification of target region in *AhCYP76AD2* covering gRNA 1 and gRNA 2 |
| CPGE_PRM00532 | tgtttcaattcttgagattcatgaga |  |
| CPGE_PRM00545 | tggtttcatctgtagaagtagcga | Amplification of target region in *AhCYP76AD5* covering gRNA 4 |
| CPGE_PRM00546 | cacaaggagatgattaatttcactca |  |
| CPGE_PRM00037 | cagctcgtgcagacctacaac | Genotyping for the presence of the *Cas9* gene in the genome |
| CPGE_PRM00038 | tgccttctaaggatagcgtg |  |

**Supplemental Table 2:** Comparison of the edit frequencies obtained from subcloning and deconvolution. It is noteworthy that subcloning demonstrates higher quality; however, it is not suitable for accurately determining edit frequencies due to the limited number of samples. The frequency of mutations was calculated as the percentage of samples with edits out of all sequenced samples, or it refers to the inferred rate of these edits in the deconvoluted Sanger sequencing chromatograms.

| **target site_callus number** | **N subcloned analyzed by Sanger sequencing** | **Frequency and modification(s) detected in subcloned fragments** | **Results of chromatogram deconvolution using ICE** |
| --- | --- | --- | --- |
| gRNA 4_5 | 8 | 25% 1 bp insertion; 12.5% 35 bp deletion | 7% 1 bp insertion;  1% 2 bp deletion |
| gRNA 4_12 | 8 | 37.5% 1 bp insertion | 9% 1 bp insertion |
| gRNA 4_17 | 8 | 25% 1 bp deletion | 15% 1 bp insertion |
| gRNA 4_25 | 8 | 25% 35 bp deletion | 8% 35 bp deletion;  7% 1 bp insertion |
| gRNA 4_52 | 8 | 12.5% 1 bp deletion; 12.5% 2 bp deletion; 12.5% 1 bp insertion | 11% 1 bp deletion;  5% 1bp insertion;  1% 2 bp deletion;  1 % 16 bp deletion |
| gRNA 4_64 | 6 | 0% efficiency | 0% efficiency |
| gRNA 1_5 | 8 | 25% 1 bp insertion | 19% 1 bp insertion |
| gRNA 1_12 | 6 | 0% efficiency | 5% 1 bp insertion;  3% 18 bp deletion |
| gRNA 1_24 | 6 | 0% efficiency | 4% 7 bp deletion;  1% 7 bp deletion |
| gRNA 1_25 | 7 | 57% 38 bp deletion | 25% 38 bp deletion: 2% 1 bp insertion;  1% 6 bp deletion;  1% 5 bp deletion |
| gRNA 1_64 | 5 | 20% 1 bp insertion | 19% 1 bp insertion;  8% 1 bp deletion |

**Supplemental Appendix 1:** DNA sequence of CPGE_VEC00447.
>CPGE_VEC00447 (17.946 bp) CGTGTCTACATTCACGTCCAAATGGGGGCTTAGATGAGAAACTTCACGATCGGCTCTAGATCGGGCCAACATGGTGGAGCACGACACTCTCGTCTACTCCAAGAATATCAAAGATACAGTCTCAGAAGACCAAAGGGCTATTGAGACTTTTCAACAAAGGGTAATATCGGGAAACCTCCTCGGATTCCATTGCCCAGCTATCTGTCACTTCATCAAAAGGACAGTAGAAAAGGAAGGTGGCACCTACAAATGCCATCATTGCGATAAAGGAAAGGCTATCGTTCAAGATGCCTCTGCCGACAGTGGTCCCAAAGATGGACCCCCACCCACGAGGAGCATCGTGGAAAAAGAAGACGTTCCAACCACGTCTTCAAAGCAAGTGGATTGATGTGATAACATGGTGGAGCACGACACTCTCGTCTACTCCAAGAATATCAAAGATACAGTCTCAGAAGACCAAAGGGCTATTGAGACTTTTCAACAAAGGGTAATATCGGGAAACCTCCTCGGATTCCATTGCCCAGCTATCTGTCACTTCATCAAAAGGACAGTAGAAAAGGAAGGTGGCACCTACAAATGCCATCATTGCGATAAAGGAAAGGCTATCGTTCAAGATGCCTCTGCCGACAGTGGTCCCAAAGATGGACCCCCACCCACGAGGAGCATCGTGGAAAAAGAAGACGTTCCAACCACGTCTTCAAAGCAAGTGGATTGATGTGATATCTCCACTGACGTAAGGGATGACGCACAATCCCACTATCCTTCGCAAGACCTTCCTCTATATAAGGAAGTTCATTTCATTTGGAGAGGACACGCTGAAATCACCAGTCTCTCTCTACAAATCTATCTCTCTCGAGGGCCCCGGGGGGCAATAAGATATGAAAAAGCCTGAACTCACCGCGACGTCTGTCGAGAAGTTTCTGATCGAAAAGTTCGACAGCGTCTCCGACCTGATGCAGCTCTCGGAGGGCGAAGAATCTCGTGCTTTCAGCTTCGATGTAGGAGGGCGTGGATATGTCCTGCGGGTAAATAGCTGCGCCGATGGTTTCTACAAAGATCGTTATGTTTATCGGCACTTTGCATCGGCCGCGCTCCCGATTCCGGAAGTGCTTGACATTGGGGAATTCAGCGAGAGCCTGACCTATTGCATCTCCCGCCGTGCACAGGGTGTCACGTTGCAAGACCTGCCTGAAACCGAACTGCCCGCTGTTCTGCAGCCGGTCGCGGAGGCCATGGATGCGATCGCTGCGGCCGATCTTAGCCAGACGAGCGGGTTCGGCCCATTCGGACCGCAAGGTAAGTTTCTGCTTCTACCTTTGATATATATATAATAATTATCATTAATTAGTAGTAATATAATATTTCAAATATTTTTTTCAAAATAAAAGAATGTAGTATATAGCAATTGCTTTTCTGTAGTTTATAAGTGTGTATATTTTAATTTATAACTTTTCTAATATATGACCAAAATTTGTTGATGTGCAGGTATCGGTCAATACACTACATGGCGTGATTTCATATGCGCGATTGCTGATCCCCATGTGTATCACTGGCAAACTGTGATGGACGACACCGTCAGTGCGTCCGTCGCGCAGGCTCTCGATGAGCTGATGCTTTGGGCCGAGGACTGCCCCGAAGTCCGGCACCTCGTGCACGCGGATTTCGGCTCCAACAATGTCCTGACGGACAATGGCCGCATAACAGCGGTCATTGACTGGAGCGAGGCGATGTTCGGGGATTCCCAATACGAGGTCGCCAACATCTTCTTCTGGAGGCCGTGGTTGGCTTGTATGGAGCAGCAGACGCGCTACTTCGAGCGGAGGCATCCGGAGCTTGCAGGATCGCCGCGGCTCCGGGCGTATATGCTCCGCATTGGTCTTGACCAACTCTATCAGAGCTTGGTTGACGGCAATTTCGATGATGCAGCTTGGGCGCAGGGTCGATGCGACGCAATCGTCCGATCCGGAGCCGGGACTGTCGGGCGTACACAAATCGCCCGCAGAAGCGCGGCCGTCTGGACCGATGGCTGTGTAGAAGTACTCGCCGATAGTGGAAACCGACGCCCCAGCACTCGTCCGAGGGCAAAGGAATAGAGTAGATGCCGACGCGTTCGAGTATTATGGCATTGGGAAAACTGTTTTTCTTGTACCATTTGTTGTGCTTGTAATTTACTGTGTTTTTTATTCGGTTTTCGCTATCGAACTGTGAAATGGAAATGGATGGAGAAGAGTTAATGAATGATATGGTCCTTTTGTTCATTCTCAAATTAATATTATTTGTTTTTTCTCTTATTTGTTGTGTGTTGAATTTGAAATTATAAGAGATATGCAAACATTTTGTTTTGAGTAAAAATGTGTCAAATCGTGGCCTCTAATGACCGAAGTTAATATGAGGAGTAAAACACTGAAGCCTGCAGGCATGCAAGCTGATCCACTAGAGGCCATGGcggccacggaattacgccaagcttgcatgcaggcctctgcagtcgacgggcccgggatccgatatctagatgcattcgcgaggtaccgagctcgaattcactggccgtttcgtctactgagcgtaaaaaccggtcccgatctagtaacatagatgacaccgcgcgcgataatttatcctagtttgcgcgctatattttgttttctatcgcgtattaaatgtataattgcgggactctaatcataaaaacccatctcataaataacgtcatgcattacatgttaattattacatgcttaacgtaattcaacagaaattatatgataatcatcgcaagaccggcaacaggattcaatcttaagaaactttattgccaaatgtttgaacgatcgagctcggggaaattcggatccccaatacttgtatggaggcctgagccgtacgggtcactggattttggttttaggaattagaaattttattgatagaagtattttacaaatacaaatacatactaagggtttcttatatgctcaacacatgagcgaaaccctataagaaccctaattcccttatctgggaactactcacacattattctggagaaaaatagagagagatagatttgtagagagagactggtgatttttgcggactctagcggtcggcatctactgcggccgcacctcaaaccttcctcttcttcttaggatcagcccttgaatcaccaccgagctgtgagagatcgatcctagtctcgtagagtccagtgatagactgatggatgagggtagcatcgagcacttctttggtagaggtgtatctcttcctatcgatggttgtatcgaagtacttgaaagcagcaggagcaccgaggttggtaagggtgaagagatggatgatgttctctgcctgttccctgataggcttatctctgtgcttgttgtaagcagacaacaccttatcgaggtttgcatcagcgaggatcacccttttagagaactcagagatctgctcgatgatctcatccaagtagtgcttgtgctgctcaacgaaaagttgcttctgctcgttatcttctggagatcccttcaacttctcgtagtgagaagcgaggtaaagaaagttaacgtacttagatgggagagcaagctcgtttcccttttgaagctcaccagcagaagcgagcatcctctttctaccgttctcgagttcgaagagtgagtactttgggagcttgatgatgagatccttcttaacctctttgtatcccttagcctcgaggaaatcgattgggttcttctcgaaagatgacctttccatgatagtgattccgagaagttccttaacagacttgagcttcttactctttcccttctcaaccttagccacaacgagaacagagtaagccacggtaggagaatcgaaaccaccgtatttcttagggtcccaatccttcttcctagcaatgagcttatcagagttcctcttagggaggatagactctttagagaatccaccggtctgcacctcggttttcttaacgatgttcacctgtggcatagagagcacctttctaacggtagcgaaatcccttcccttatcccacacgatctcacctgtttcaccgtttgtctcgatgagtggcctctttctgatctcaccgttagcgagggtaatctcggtcttgaagaaattcatgatgttagagtagaagaaatacttagcggtagcctttccgatctcttgctcagacttagcgatcatcttcctcacatcgtacaccttgtaatcaccgtacacgaactctgactcgagcttaggatacttcttgatgagagcggttccaacaacagcgttaaggtaagcatcgtgagcgtggtggtagttgttgatttccctcaccttgtagaattggaaatcctttctgaaatcagacacgagctttgacttgagggtgataaccttcacttccctgatcaacttatcgttctcatcgtacttggtgttcatcctagaatcgaggatctgtgcaacgtgcttagtgatctgcctggtttccacaagctgcctcttgatgaatcctgccttatccaattcagagagtcctcccctctcagccttagtcaagttatcgaactttctctgagtgatgagcttagcgttgaggagctgcctccaatagttcttcattttcttcacaacctcttcacttggcacgttatcactcttacccctgttcttatcagacctggtgagcaccttgttatcgatagaatcatccttcaagaatgactgtggcacgatatgatcaacatcgtaatcagagagcctgttgatatccaactcttgatccacatacatatcccttccgttctggaggtagtagaggtagagcttctcattctggagctgagtgttctcaacagggtgctctttgaggatctgagatccaagctctttgataccttcctcgatcctcttcatcctttccctagagttcttctgtcccttctgagtggtctggttctctctagccatttcgatcacgatgttctcaggcttatgccttcccatcaccttcaccaactcatccacaaccttcacagtctggaggattcccttcttgattgcaggagatccagcgaggttagcgatatgctcatggagactatcaccctgtcctgaaacctgagccttctggatatcctctttaaaggtgagagaatcatcgtggatgagctgcatgaagtttctgttagcgaatccatcagacttgaggaaatcaaggattgtctttccagactgcttatccctgattccgttaatgagctttcttgagagccttccccaaccagtgtatcttcttctcttcaactgcttcatcaccttatcatcgaagagatgagcgtaggtcttgagcctttcttcaatcatctctctatcttcaaagagggtgagggtaagaacgatatcctccaagatatcctcgttttcctcgttatccaagaaatccttatccttaatgatcttgaggagatcgtggtaggttccgagagatgcgttgaacctatcctcaacaccagaaatctcaactgaatcgaagcactcgattttcttgaagtaatcctctttgagctgcttcacggtcacctttctgttggtcttgaacaagagatcaacgatagccttcttttgctcacctgacaaaaaagcaggcttcctcattccctcggtcacgtacttaaccttggtcaactcgttgtacacggtgaagtactcgtagagcaaagagtgcttagggagcaccttctcgtttggaaggttcttatcgaagttggtcatcctctcgatgaaagactgagcactagcacccttatccaccacctcttcgaagttccaaggggtgatggtttcctcagactttctggtcatccaagcgaatcttgagtttcctctagcgagaggtcccacgtagtaagggattctgaaggtgagaatcttctcaatcttttccctgttatccttgaggaatgggtagaaatcctcttgccttctaaggatagcgtgcaactctccgaggtggatctgatgagggatagatccgttatcgaaggtcctctgctttctgagaagatcctctctattgagcttcacgaggagttcctcggttccatccatcttctcgaggataggcttgatgaacttgtagaactcttcttgagatgcaccaccatcgatgtaaccagcgtatccgttcttagactgatcgaagaaaatctctttgtacttctctgggagctgctgtctaacaagagccttgagaagtgtgagatcctggtggtgctcatcgtatctcttgatcatagaagctgagagtggagccttggtgatctcggtgttcactctgaggatatcactgaggaggatagcatcagagaggttcttagcagcgaggaacaaatcagcgtactgatctccgatctgagcgaggaggttatcgagatcatcatcgtaggtatcctttgagagctggagctttgcatcctcagcgagatcgaagttagacttgaagttaggggtgagtccgagagagagagcgatcaagtttccgaaaagtccgttcttcttctcaccagggagctgagcaatgaggttctcaagccttcttgacttagagagcctagcagagaggatagccttagcatccacacctgaagcgttgatagggttctcttcgaaaagctggttgtaggtctgcacgagctggatgaacaacttatccacatcagagttatcagggttgagatcaccctcgatgaggaagtgtcctctgaacttgatcatgtgagcgagagcgaggtagatgagcctgagatcagccttatcagtagaatcaacgagcttctttctgaggtggtagatagtagggtacttctcgtggtatgccacctcatcaacgatgtttccgaagatagggtgcctctcgtgcttcttatcttcttccacgaggaatgactcttcgagcctgtggaagaatgaatcatccactttagccatctcgttagagaagatctcttggaggtagcagatcctgttctttcttctggtgtaccttcttctagcggttctcttgagtctggtagcctcagcagtttcaccagaatcgaagaggagagcaccgataaggtttttcttgatagagtgcctatcggtgtttccgagaaccttgaacttcttagatggcaccttgtactcatcggtgatcacagcccatcccacagagttagttccgatatcgagtccgatagagtacttcttatccatgctgcacatacataacatatcaagatcagaacacacatatacacacacaaatacaatcaagtcaacaactccaaaaagtccagatctacatatatacatacgtaaataacaaaatcatgtaaataatcacaatcatgtaatccagatctatgcacatatatatatacacaattaataaaaaaaatgatataacagatctatatctatgtatgtaacaacacaatcagatgagagaagtgatgttttcagatctgtatacatacaaacacaaacagatgaacaattgatacgtagatccatatgtatacgtacaattagctacacgattaaatgaaaaaaatcaacgatttcggattggtacacacaaacgcaacaatatgaagaaattcatatctgattagatataaacataaccacgtgtagatacacagtcaaatcaacaaatttatagcttctaaacggatgagatgaacaagataaagatattcacataaggcatacataagataagcagattaacaaactagcaataatacatacctaattaaaacaaggaataacagagagagagagagagagagagatttaccttgaaaatgaagaggagaagagaggatttcttaaaattgggggtagagaaagaaagatgatgaattgtgagaaaggagagatagaagggggggttgtatatataggctgtagaagattatttttgtgtttgaggcggtgaaggaagaggggatctgactatgacacgtttgcggttacgtatttcgataggagtctttcaacgcttaacgccgttactctatatgaccgtttgggccgtaacggggccgtttgttaacgctgatgttgattcttttctttctttctttcttccttttttaaagaagcaattgtacaatcgttgctagctgtcaaacggataattcggatacggatatgcctatattcatatccgtaatttttcaatctacgctggtctactgagtctaatgccaactttgtacaagaaagctgggtctagaaaaaaagcaccgactcggtgccactttttcaagttgataacggactagccttattttaacttgctatttctagctctaaaacgctctgactgaatctggaatCAATcactacttcgactctagctgtatataaactcagcttcgttttcttatctaagcgatgtgggacttttgaagattgttttcaacttaaatgggcctatataagaaatactattgttctttcccatataaatgggcctgcttctcttctttcagattcccaggggccttttgaagattatcttcatatcttaagaatgaagatgttttattcaatcaaattcttgaaggttcgatgcctaatcattctaatcctgggacaaactatgaaacaagatacaaaaactccgaatggaaagttaaaaagaagaaaacgaaagctacggttcaagaaaatgtaagctgataaacaaaaaaaaactgtatgaacgaagaagaagaaaaaaagaccgtaatgccaactttgtacaagaaagctgggtctagaaaaaaagcaccgactcggtgccactttttcaagttgataacggactagccttattttaacttgctatttctagctctaaaacccgaccgtgttcttgctcccCAATcactacttcgactctagctgtatataaactcagcttcgttttcttatctaagcgatgtgggacttttgaagattgttttcaacttaaatgggcctatataagaaatactattgttctttcccatataaatgggcctgcttctcttctttcagattcccaggggccttttgaagattatcttcatatcttaagaatgaagatgttttattcaatcaaattcttgaaggttcgatgcctaatcattctaatcctgggacaaactatgaaacaagatacaaaaactccgaatggaaagttaaaaagaagaaaacgaaagctacggttcaagaaaatgtaagctgataaacaaaaaaaaactgtatgaacgaagaagaagaaaaaaagagggtaatgccaactttgtacaagaaagctgggtctagaaaaaaagcaccgactcggtgccactttttcaagttgataacggactagccttattttaacttgctatttctagctctaaaaccggccctttaatttccttatCAATcactacttcgactctagctgtatataaactcagcttcgttttcttatctaagcgatgtgggacttttgaagattgttttcaacttaaatgggcctatataagaaatactattgttctttcccatataaatgggcctgcttctcttctttcagattcccaggggccttttgaagattatcttcatatcttaagaatgaagatgttttattcaatcaaattcttgaaggttcgatgcctaatcattctaatcctgggacaaactatgaaacaagatacaaaaactccgaatggaaagttaaaaagaagaaaacgaaagctacggttcaagaaaatgtaagctgataaacaaaaaaaaactgtatgaacgaagaagaagaaaaaaagaagctaatgccaactttgtacaagaaagctgggtctagaaaaaaagcaccgactcggtgccactttttcaagttgataacggactagccttattttaacttgctatttctagctctaaaactggcatgtctaagggattgcCAATcactacttcgactctagctgtatataaactcagcttcgttttcttatctaagcgatgtgggacttttgaagattgttttcaacttaaatgggcctatataagaaatactattgttctttcccatataaatgggcctgcttctcttctttcagattcccaggggccttttgaagattatcttcatatcttaagaatgaagatgttttattcaatcaaattcttgaaggttcgatgcctaatcattctaatcctgggacaaactatgaaacaagatacaaaaactccgaatggaaagttaaaaagaagaaaacgaaagctacggttcaagaaaatgtaagctgataaacaaaaaaaaactgtatgaacgaagaagaagaaaaaaaggcaatctactgacctaggccttaaGGGCCAGATCTTGGGCCCGGTACCCGATCAGATTGTCGTTTCCCGCCTTCGGTTTAAACTATCAGTGTTTGACAGGATATATTGGCGGGTAAACCTAAGAGAAAAGAGCGTTTATTAGAATAATCGGATATTTAAAAGGGCGTGAAAAGGTTTATCCGTTCGTCCATTTGTATGTGCATGCCAACCACAGGGTTCCCCTCGGGAGTGCTTGGCATTCCGTGCGATAATGACTTCTGTTCAACCACCCAAACGTCGGAAAGCCTGACGACGGAGCAGCATTCCAAAAAGATCCCTTGGCTCGTCTGGGTCGGCTAGAAGGTCGAGTGGGCTGCTGTGGCTTGATCCCTCAACGCGGTCGCGGACGTAGCGCAGCGCCGAAAAATCCTCGATCGCAAATCCGACGCTGTCGAAAAGCGTGATCTGCTTGTCGCTCTTTCGGCCGACGTCCTGGCCAGTCATCACGCGCCAAAGTTCCGTCACAGGATGATCTGGCGCGAGTTGCTGGATCTCGCCTTCAATCCGGGTCTGTGGCGGGAACTCCACGAAAATATCCGAACGCAGCAAGATATCGCGGTGCATCTCGGTCTTGCCTGGGCAGTCGCCGCCGACGCCGTTGATGTGGACGCCGAAAAGGATCTAGGTGAAGATCCTTTTTGATAATCTCATGACCAAAATCCCTTAACGTGAGTTTTCGTTCCACTGAGCGTCAGACCCCGTAGAAAAGATCAAAGGATCTTCTTGAGATCCTTTTTTTCTGCGCGTAATCTGCTGCTTGCAAACAAAAAAACCACCGCTACCAGCGGTGGTTTGTTTGCCGGATCAAGAGCTACCAACTCTTTTTCCGAAGGTAACTGGCTTCAGCAGAGCGCAGATACCAAATACTGTTCTTCTAGTGTAGCCGTAGTTAGGCCACCACTTCAAGAACTCTGTAGCACCGCCTACATACCTCGCTCTGCTAATCCTGTTACCAGTGGCTGCTGCCAGTGGCGATAAGTCGTGTCTTACCGGGTTGGACTCAAGACGATAGTTACCGGATAAGGCGCAGCGGTCGGGCTGAACGGGGGGTTCGTGCACACAGCCCAGCTTGGAGCGAACGACCTACACCGAACTGAGATACCTACAGCGTGAGCTATGAGAAAGCGCCACGCTTCCCGAAGGGAGAAAGGCGGACAGGTATCCGGTAAGCGGCAGGGTCGGAACAGGAGAGCGCACGAGGGAGCTTCCAGGGGGAAACGCCTGGTATCTTTATAGTCCTGTCGGGTTTCGCCACCTCTGACTTGAGCGTCGATTTTTGTGATGCTCGTCAGGGGGGCGGAGCCTATGGAAAAACGCCAGCAACGCGGCCTTTTTACGGTTCCTGGCCTTTTGCTGGCCTTTTGCTCACATGTTCTTTCCTGCGTTATCCCCTGATTCTGTGGATAACCGATTACCGCCTTTGAGTGAGCTGATACCGCTCGCCGCAGCCGAACGACCGAGCGCAGCGAGTCAGTGAGCGAGGAAGCGGAAGAGCGCCTGATGCGGTATTTTCTCCTTACGCATCTGTGCGGTATTTCACACCGCATATGGTGCACTCTCAGTACAATCTGCTCTGATGCCGCATAGTTAAGCCAGTATACACTCCGCTATCGCTACGTGACTGGGTCATGGCTGCGCCCCGACACCCGCCAACACCCGCTGACGCGCCCTGACGGGCTTGTCTGCTCCCGGCATCCGCTTACAGACAAGCTGTGACCGTCTCCGGGAGCTGCATGTGTCAGAGGTTTTCACCGTCATCACCGAAACGCGCGAGGCAGGGGTACGTCGAGGTCGATCCAACCCCTCCGCTGCTATAGTGCAGTCGGCTTCTGACGTTCAGTGCAGCCGTCTTCTGAAAACGACATGTCGCACAAGTCCTAAGTTACGCGACAGGCTGCCGCCCTGCCCTTTTCCTGGCGTTTTCTTGTCGCGTGTTTTAGTCGCATAAAGTAGAATACTTGCGACTAGAACCGGAGACATTACGCCATGAACAAGAGCGCCGCCGCTGGCCTGCTGGGCTATGCCCGCGTCAGCACCGACGACCAGGACTTGACCAACCAACGGGCCGAACTGCACGCGGCCGGCTGCACCAAGCTGTTTTCCGAGAAGATCACCGGCACCAGGCGCGACCGCCCGGAGCTGGCCAGGATGCTTGACCACCTACGCCCTGGCGACGTTGTGACAGTGACCAGGCTAGACCGCCTGGCCCGCAGCACCCGCGACCTACTGGACATTGCCGAGCGCATCCAGGAGGCCGGCGCGGGCCTGCGTAGCCTGGCAGAGCCGTGGGCCGACACCACCACGCCGGCCGGCCGCATGGTGTTGACCGTGTTCGCCGGCATTGCCGAGTTCGAGCGTTCCCTAATCATCGACCGCACCCGGAGCGGGCGCGAGGCCGCCAAGGCGCGAGGCGTGAAGTTTGGCCCCCGCCCTACCCTCACCCCGGCACAGATCGCGCACGCCCGCGAGCTGATCGACCAGGAAGGCCGCACCGTGAAAGAGGCGGCTGCACTGCTTGGCGTGCATCGCTCGACCCTGTACCGCGCACTTGAGCGCAGCGAGGAAGTGACGCCCACCGAGGCCAGGCGGCGCGGTGCCTTCCGTGAGGACGCATTGACCGAGGCCGACGCCCTGGCGGCCGCCGAGAATGAACGCCAAGAGGAACAAGCATGAAACCGCACCAGGACGGCCAGGACGAACCGTTTTTCATTACCGAAGAGATCGAGGCGGAGATGATCGCGGCCGGGTACGTGTTCGAGCCGCCCGCGCACGTCTCAACCGTGCGGCTGCATGAAATCCTGGCCGGTTTGTCTGATGCCAAGCTCGCGGCCTGGCCGGCGAGCTTGGCCGCTGAAGAAACCGAGCGCCGCCGTCTAAAAAGGTGATGTGTATTTGAGTAAAACAGCTTGCGTCATGCGGTCGCTGCGTATATGATGCGATGAGTAAATAAACAAATACGCAAGGGGAACGCATGAAGGTTATCGCTGTACTTAACCAGAAAGGCGGGTCAGGCAAGACGACCATCGCAACCCATCTAGCCCGCGCCCTGCAACTCGCCGGGGCCGATGTTCTGTTAGTCGATTCCGATCCCCAGGGCAGTGCCCGCGATTGGGCGGCCGTGCGGGAAGATCAACCGCTAACCGTTGTCGGCATCGACCGCCCGACGATTGACCGCGACGTGAAGGCCATCGGCCGGCGCGACTTCGTAGTGATCGACGGAGCGCCCCAGGCGGCGGACTTGGCTGTGTCCGCGATCAAGGCAGCCGACTTCGTGCTGATTCCGGTGCAGCCAAGCCCTTACGACATATGGGCCACCGCCGACCTGGTGGAGCTGGTTAAGCAGCGCATTGAGGTCACGGATGGAAGGCTACAAGCGGCCTTTGTCGTGTCGCGGGCGATCAAAGGCACGCGCATCGGCGGTGAGGTTGCCGAGGCGCTGGCCGGGTACGAGCTGCCCATTCTTGAGTCCCGTATCACGCAGCGCGTGAGCTACCCAGGCACTGCCGCCGCCGGCACAACCGTTCTTGAATCAGAACCCGAGGGCGACGCTGCCCGCGAGGTCCAGGCGCTGGCCGCTGAAATTAAATCAAAACTCATTTGAGTTAATGAGGTAAAGAGAAAATGAGCAAAAGCACAAACACGCTAAGTGCCGGCCGTCCGAGCGCACGCAGCAGCAAGGCTGCAACGTTGGCCAGCCTGGCAGACACGCCAGCCATGAAGCGGGTCAACTTTCAGTTGCCGGCGGAGGATCACACCAAGCTGAAGATGTACGCGGTACGCCAAGGCAAGACCATTACCGAGCTGCTATCTGAATACATCGCGCAGCTACCAGAGTAAATGAGCAAATGAATAAATGAGTAGATGAATTTTAGCGGCTAAAGGAGGCGGCATGGAAAATCAAGAACAACCAGGCACCGACGCCGTGGAATGCCCCATGTGTGGAGGAACGGGCGGTTGGCCAGGCGTAAGCGGCTGGGTTGTCTGCCGGCCCTGCAATGGCACTGGAACCCCCAAGCCCGAGGAATCGGCGTGAGCGGTCGCAAACCATCCGGCCCGGTACAAATCGGCGCGGCGCTGGGTGATGACCTGGTGGAGAAGTTGAAGGCGGCGCAGGCCGCCCAGCGGCAACGCATCGAGGCAGAAGCACGCCCCGGTGAATCGTGGCAAGCGGCCGCTGATCGAATCCGCAAAGAATCCCGGCAACCGCCGGCAGCCGGTGCGCCGTCGATTAGGAAGCCGCCCAAGGGCGACGAGCAACCAGATTTTTTCGTTCCGATGCTCTATGACGTGGGCACCCGCGATAGTCGCAGCATCATGGACGTGGCCGTTTTCCGTCTGTCGAAGCGTGACCGACGAGCTGGCGAGGTGATCCGCTACGAGCTTCCAGACGGGCACGTAGAGGTTTCCGCAGGGCCGGCCGGCATGGCGAGTGTGTGGGATTACGACCTGGTACTGATGGCGGTTTCCCATCTAACCGAATCCATGAACCGATACCGGGAAGGGAAGGGAGACAAGCCCGGCCGCGTGTTCCGTCCACACGTTGCGGACGTACTCAAGTTCTGCCGGCGAGCCGATGGCGGAAAGCAGAAAGACGACCTGGTAGAAACCTGCATTCGGTTAAACACCACGCACGTTGCCATGCAGCGTACGAAGAAGGCCAAGAACGGCCGCCTGGTGACGGTATCCGAGGGTGAAGCCTTGATTAGCCGCTACAAGATCGTAAAGAGCGAAACCGGGCGGCCGGAGTACATCGAGATCGAGCTAGCTGATTGGATGTACCGCGAGATCACAGAAGGCAAGAACCCGGACGTGCTGACGGTTCACCCCGATTACTTTTTGATCGATCCCGGCATCGGCCGTTTTCTCTACCGCCTGGCACGCCGCGCCGCAGGCAAGGCAGAAGCCAGATGGTTGTTCAAGACGATCTACGAACGCAGTGGCAGCGCCGGAGAGTTCAAGAAGTTCTGTTTCACCGTGCGCAAGCTGATCGGGTCAAATGACCTGCCGGAGTACGATTTGAAGGAGGAGGCGGGGCAGGCTGGCCCGATCCTAGTCATGCGCTACCGCAACCTGATCGAGGGCGAAGCATCCGCCGGTTCCTAATGTACGGAGCAGATGCTAGGGCAAATTGCCCTAGCAGGGGAAAAAGGTCGAAAAGGTCTCTTTCCTGTGGATAGCACGTACATTGGGAACCCAAAGCCGTACATTGGGAACCGGAACCCGTACATTGGGAACCCAAAGCCGTACATTGGGAACCGGTCACACATGTAAGTGACTGATATAAAAGAGAAAAAAGGCGATTTTTCCGCCTAAAACTCTTTAAAACTTATTAAAACTCTTAAAACCCGCCTGGCCTGTGCATAACTGTCTGGCCAGCGCACAGCCGAAGAGCTGCAAAAAGCGCCTACCCTTCGGTCGCTGCGCTCCCTACGCCCCGCCGCTTCGCGTCGGCCTATCGCGGCCGCTGGCCGCTCAAAAATGGCTGGCCTACGGCCAGGCAATCTACCAGGGCGCGGACAAGCCGCGCCGTCGCCACTCGACCGCCGGCGCCCACATCAAGGCACCGGTGGGTATGCCTGACGATGCGTGGAGACCGAAACCTTGCGCTCGTTCGCCAGCCAGGACAGAAATGCCTCGACTTCGCTGCTGCCCAAGGTTGCCGGGTGACGCACACCGTGGAAACGGATGAAGGCACGAACCCAGTGGACATAAGCCTGTTCGGTTCGTAAGCTGTAATGCAAGTAGCGTATGCGCTCACGCAACTGGTCCAGAACCTTGACCGAACGCAGCGGTGGTAACGGCGCAGTGGCGGTTTTCATGGCTTGTTATGACTGTTTTTTTGGGGTACAGTCTATGCCTCGGGCATCCAAGCAGCAAGCGCGTTACGCCGTGGGTCGATGTTTGATGTTATGGAGCAGCAACGATGTTACGCAGCAGGGCAGTCGCCCTAAAACAAAGTTAAACATCATGAGGGAAGCGGTGATCGCCGAAGTATCGACTCAACTATCAGAGGTAGTTGGCGTCATCGAGCGCCATCTCGAACCGACGTTGCTGGCCGTACATTTGTACGGCTCCGCAGTGGATGGCGGCCTGAAGCCACACAGTGATATTGATTTGCTGGTTACGGTGACCGTAAGGCTTGATGAAACAACGCGGCGAGCTTTGATCAACGACCTTTTGGAAACTTCGGCTTCCCCTGGAGAGAGCGAGATTCTCCGCGCTGTAGAAGTCACCATTGTTGTGCACGACGACATCATTCCGTGGCGTTATCCAGCTAAGCGCGAACTGCAATTTGGAGAATGGCAGCGCAATGACATTCTTGCAGGTATCTTCGAGCCAGCCACGATCGACATTGATCTGGCTATCTTGCTGACAAAAGCAAGAGAACATAGCGTTGCCTTGGTAGGTCCAGCGGCGGAGGAACTCTTTGATCCGGTTCCTGAACAGGATCTATTTGAGGCGCTAAATGAAACCTTAACGCTATGGAACTCGCCGCCCGACTGGGCTGGCGATGAGCGAAATGTAGTGCTTACGTTGTCCCGCATTTGGTACAGCGCAGTAACCGGCAAAATCGCGCCGAAGGATGTCGCTGCCGACTGGGCAATGGAGCGCCTGCCGGCCCAGTATCAGCCCGTCATACTTGAAGCTAGACAGGCTTATCTTGGACAAGAAGAAGATCGCTTGGCCTCGCGCGCAGATCAGTTGGAAGAATTTGTCCACTACGTGAAAGGCGAGATCACCAAGGTAGTCGGCAAATAATGTCTAACAATTCGTTCAAGCCGACGCCGCTTCGCGGCGCGGCTTAACTCAAGCGTTAGATGCACTAAGCACATAATTGCTCACAGCCAAACTATCAGGTCAAGTCTGCTTTTATTATTTTTAAGCGTGCATAATAAGCCCTACACAAATTGGGAGATATATCATGAAAGGCTGGCTTTTTCTTGTTATCGCAATAGTTGGCGAAGTAATCGCAACATAGCTTGCTTGGTCGTTCCGCGTGAACGTCGGCTCGATTGTACCTGCGTTCAAATACTTTGCGATCGTGTTGCGCGCCTGCCCGGTGCGTCGGCTGATCTCACGGATCGACTGCTTCTCTCGCAACGCCATCCGACGGATGATGTTTAAAAGTCCCATGTGGATCACTCCGTTGCCCCGTCGCTCACCGTGTTGGGGGGAAGGTGCACATGGCTCAGTTCTCAATGGAAATTATCTGCCTAACCGGCTCAGTTCTGCGTAGAAACCAACATGCAAGCTCCACCGGGTGCAAAGCGGCAGCGGCGGCAGGATATATTCAATTGTAAATGGCTTCATGTCCGGGAAATCTACATGGATCAGCAATGAGTATGATGGTCAATATGGAGAAAAAGAAAGAGTAATTACCAATTTTTTTTCAATTCAAAAATGTAGATGTCCGCAGCGTTATTATAAAATGAAAGTACATTTTGATAAAACGACAAATTACGATCCGTCGTATTTATAGGCGAAAGCAATAAACAAATTATTCTAATTCGGAAATCTTTATTTCGA
